# Lingering single-strand breaks trigger Rad51-independent homology-directed repair of collapsed replication forks in polynucleotide kinase/phosphatase mutant of fission yeast

**DOI:** 10.1101/157354

**Authors:** Arancha Sanchez, Mariana C. Gadaleta, Oliver Limbo, Paul Russell

**Affiliations:** Department of Molecular Medicine The Scripps Research Institute La Jolla, CA, 92037

**Keywords:** polynucleotide kinase phosphatase, PNKP, single-strand break repair, DNA repair, DNA damage, replication fork collapse, genomic instability, genome integrity, homologous recombination, Schizosaccharomyces pombe, fission yeast

## Abstract

The DNA repair enzyme polynucleotide kinase/phosphatase (PNKP) protects genome integrity by restoring ligatable 5’-phosphate and 3’-hydroxyl termini at single-strand breaks (SSBs). In humans, PNKP mutations underlie the neurological disease known as MCSZ, but these individuals are not predisposed for cancer, implying effective alternative repair pathways in dividing cells. Homology-directed repair (HDR) of collapsed replication forks was proposed to repair SSBs in PNKP-deficient cells, but the critical HDR protein Rad51 is not required in PNKP-null (*pnk1Δ*) cells of *Schizosaccharomyces pombe.* Here, we report that *pnk1Δ* cells have enhanced requirements for Rad3 (ATR/Mec1) and Chk1 checkpoint kinases, and the multi-BRCT domain protein Brc1 that binds phospho-histone H2A (γH2A) at damaged replication forks. The viability of *pnk1Δ* cells depends on Mre11 and Ctp1 (CtIP/Sae2) double-strand break (DSB) resection proteins, Rad52 DNA strand annealing protein, Mus81-Eme1 Holliday junction resolvase, and Rqh1 (BLM/WRN/Sgs1) DNA helicase. Eliminating Pnk1 strongly sensitizes *mre11Δ pku80Δ* cells to DNA damaging agents that collapse replication forks, indicating a requirement for Mre11-Rad50-Nbs1 (MRN) protein complex that cannot be efficiently replaced by Exo1 5’-3’ exonuclease. Coupled with increased sister chromatid recombination and Rad52 repair foci in *pnk1Δ* cells, these findings indicate that lingering SSBs in *pnk1Δ* cells trigger Rad51-independent homology-directed repair of collapsed replication forks.

**AUTHOR SUMMARY:** DNA is constantly damaged by normal cellular metabolism, for example production of reactive oxygen species, or from exposure to external DNA damaging sources, such as radiation from the sun or chemicals in the environment. These genotoxic agents create thousands of single-strand breaks/cell/day in the human body. An essential DNA repair protein known as polynucleotide kinase/phosphatase (PNKP) makes sure the single-strand breaks have 5’ phosphate and 3’ hydroxyl ends suitable for healing by DNA ligase. Mutations that reduce PNKP activity cause a devastating neurological disease but surprisingly not cancer, suggesting that other DNA repair mechanisms step into the breach in dividing PNKP-deficient cells. One popular candidate was homology-directed repair (HDR) of replication forks that collapse at single-strand breaks, but the crucial HDR protein Rad51 was found to be non-essential in PNKP-deficient cells of fission yeast. In this study, Sanchez and Russell revive the HDR model by showing that SSBs in PNKP-deficient cells are repaired by a variant HDR mechanism that bypasses the requirement for Rad51. Notably, Mus81 endonuclease that resolves sister chromatid recombination structures formed during HDR of collapsed replication forks was found to be essential in PNKP-deficient cells.

## INTRODUCTION

Maintenance of genome integrity depends on the accurate repair of DNA lesions that sever one or both strands of the double-helix. Single-strand breaks (SSBs) are by far the most abundant DNA scission, occurring at frequencies of thousands/cell/day in proliferating human cells (1). SSBs are formed by many mechanisms, including oxidative attack of the sugar-phosphate backbone by endogenous reactive oxygen species (ROS), by base- and nucleotide excision repair, through the activity of anti-cancer drugs such as camptothecin or bleomycins, or by exposure to other DNA damaging agents. These SSBs often have 5’-hydroxyl or 3’-phosphate termini that prevent ligation. Polynucleotide kinase phosphatase (PNKP) is a bifunctional enzyme that restores 5’-phosphate and 3’-hydroxyl to these DNA ends (2, 3). PNKP’s importance is indicated by its conservation throughout eukaryotic evolution, although some species such as *Saccharomyces cerevisiae* have only retained the phosphatase domain (4).

The consequences of eliminating PNKP activity varies dramatically in eukaryotes. At one extreme, deleting the PNKP gene in mice causes early embryonic lethality (5). PNKP probably plays an equally important role in humans, as a rare autosomal recessive disease characterized by microcephaly, early-onset intractable seizures and developmental delay (denoted MCSZ) was traced to partial loss-of-function mutations in the PNKP gene (6-8). MCSZ is not associated with cancer; indeed, neurodegeneration in the absence of cancer predisposition appears to be a typical consequence of SSB repair defects in humans (9). In contrast to mammals, *S. cerevisiae* cells lacking the DNA 3’ phosphatase encoded by *TPP1* display no obvious phenotypes or sensitivity to DNA damaging agents (10). However, requirements for Tpp1 are revealed when other DNA repair pathways are inactivated. Most notably, in cells lacking the apurinic/apyrimidinic (AP) endonucleases Apn1 and Apn2, deletion of *TPP1* increases cellular sensitivity to several DNA damaging agents, including the DNA alkylating agent methyl methanesulfonate (MMS) and the topoisomerase I inhibitor camptothecin (CPT) (10, 11). These AP endonucleases process DNA ends with various 3’-terminal blocking lesions, including 3’ phosphoglycolate (3’-PG), 3′-unsaturated aldehydic, α,β-4-hydroxy-2-pentenal (3′-dRP), and 3’-phosphates. PNKP is not essential in the fission yeast *Schizosaccharomyces pombe*, but *pnk1Δ* cells are sensitive to a variety of DNA damaging agents, most notably CPT (12-14). These phenotypes were attributed to loss of Pnk1 phosphatase activity, as they are rescued by expression of *TPP1* or kinase-null mutations of *pnk1,* but not *pnk1* alleles that eliminate phosphatase activity (14). In contrast to *S. cerevisiae,* in which *tpp1Δ apn1Δ apn2Δ* cells display no obvious growth defect (10), in *S. pombe pnk1Δ apn2Δ* cells are inviable (14).

If SSBs with 5’-hydroxyl or 3’-phosphate are left unrepaired in PNKP-deficient cells, progression through S-phase should lead to replication fork collapse, resulting in one-ended double-strand breaks (DSBs) (1). These DNA lesions are subject to homology-directed repair (HDR), which initiates when an endonuclease consisting of Mre11-Rad50-Nbs1 (MRN) protein complex and Ctp1 (CtIP/Sae2) binds the DSB and progressively clips the 5’ strand, generating a 3’ single-strand DNA (ssDNA) overhang (15-17). This ssDNA is coated with Replication Protein A (RPA), which is then replaced by Rad51 recombinase by a mechanism requiring Rad52 strand-annealing protein. Rad51 catalyzes the homology search and invasion of the intact sister chromatid, culminating in restoration of the replication fork. This fork repair mechanism produces a DNA joint molecule, aka Holliday junction (HJ), that must be resolved to allow chromosome segregation during mitosis. Replication-coupled single-strand break repair as outlined above has been widely proposed as an alternative mechanism for repairing SSBs in PNKP-deficient cells (1, 18, 19). However, data supporting this model are weak. Notably, Rad51 is not required in *pnk1Δ* mutants of fission yeast (14). Nor does elimination of TPP1 cause any reported phenotype in *rad52Δ* cells of budding yeast, although significant growth defects appear when AP endonucleases are also eliminated in this genetic background (10). Most critically, it is unknown whether HJ resolvases are required in PNKP-deficient cells, which is a decisive prediction of the replication-coupled single-strand break repair model.

Brc1 is a fission yeast protein with 6 BRCT (BRCA1 C-terminal) domains that is structurally related to budding yeast Rtt107 and human PTIP (20, 21). The C-terminal pair of BRCT domains in Brc1 bind phospho-histone H2A (γH2A), equivalent to mammalian γH2AX, which is formed by Tel1 (ATM) and Rad3 (ATR/Mec1) checkpoint kinases at DSBs and damaged or stalled replication forks (21-23). Brc1 is not required for DSB repair but it plays an important role in recovery from replication fork collapse (13, 22, 24-28). We recently discovered a synergistic negative genetic interaction involving *brc1Δ* and *pnk1Δ* (13), suggesting that *pnk1Δ* cells suffer increased rates of fork collapse. This result presents a conundrum, because as mentioned above, the critical HDR protein Rad51 is not required for the viability of *pnk1Δ* cells (14). Here, we investigate the genetic requirements for surviving PNKP deficiency in fission yeast, uncovering crucial roles for key HDR proteins such as Mre11, Rad52 and Mus81 in a variant mechanism of replication fork repair that does not require Rad51.

## RESULTS

### Brc1 binding to γH2A is critical in *pnk1Δ* cells

Epistatic mini-array profiling (E-MAP) screens identified synergistic negative genetic interactions involving *brc1Δ* and *pnk1Δ,* indicating that Brc1 helps to maintain cell viability when Pnk1 activity is lost (13, 29, 30). We confirmed this synthetic sick interaction in spot dilution assays in which the colony size of *brc1Δ pnk1Δ* mutants were reduced compared to either single mutant (Figure 1, untreated panels). The growth defect of *brc1Δ pnk1Δ* cells was enhanced in the presence of MMS or CPT, which produce DNA lesions that can be processed to yield SSBs with 3’ phosphate (Figure 1). Failure to repair these SSBs before entry into S-phase would be expected to increase the frequency of replication fork collapse.

**Figure 1.**
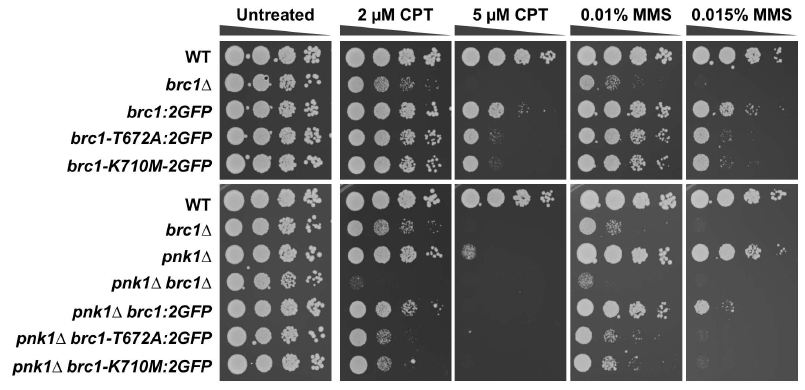
Brc1 binding to γH2A is important in *pnk1Δ* cells. Tenfold serial dilutions of cells were exposed to the indicated DNA damaging agents. Plates were incubated at 30°C for 3 to 4 days. Note that the *brc1-T672A* and *brc1-K710M* alleles contain a C-terminal 2GFP tag, which under some conditions can be observed to partially impair Brc1, thus strains with these alleles should be compared to wild type Brc1 tagged with 2GFP *(brc1:2GFP).*

Brc1 is thought to act as a scaffold protein to promote replication fork stability and repair (21, 28). These activities partially depend on the ability of Brc1 to bind γH2A through its C-terminal pair of BRCT domains. The crystal structure of these domains bound to γH2A peptide allowed us to design T672A and K710M mutations that specifically disrupt the γH2A-binding pocket in Brc1 (21). These mutations did not cause an obvious growth defect in the *pnk1Δ* background but they strongly enhanced sensitivity to MMS or CPT (Figure 1). From these data, we conclude that Brc1 binding to γH2A is critical when *pnk1Δ* cells are treated with genotoxins that causes formation of SSBs with 3’ phosphate.

### ATR/Rad3 and Chk1 checkpoint kinases are crucial in *pnk1Δ* cells

The requirement for Brc1 binding to γH2A in *pnk1Δ* cells suggested that unrepaired SSBs in these cells triggers a DNA damage response involving the master checkpoint kinase ATR, known as Rad3 in fission yeast (31). Indeed, *pnk1Δ rad3Δ* colony size was reduced compared to either single mutant (Figure 2A, untreated panel). This negative genetic interaction became more obvious when *pnk1Δ rad3Δ* cells were grown in the presence of CPT, MMS or the replication inhibitor hydroxyurea (HU) (Figure 2A). Elimination of Brc1 further impaired growth in *pnk1Δ rad3Δ* cells (Figure 2A), which is consistent with previous studies indicating that Brc1 has both Rad3-dependent and independent activities (13).

**Figure 2.**
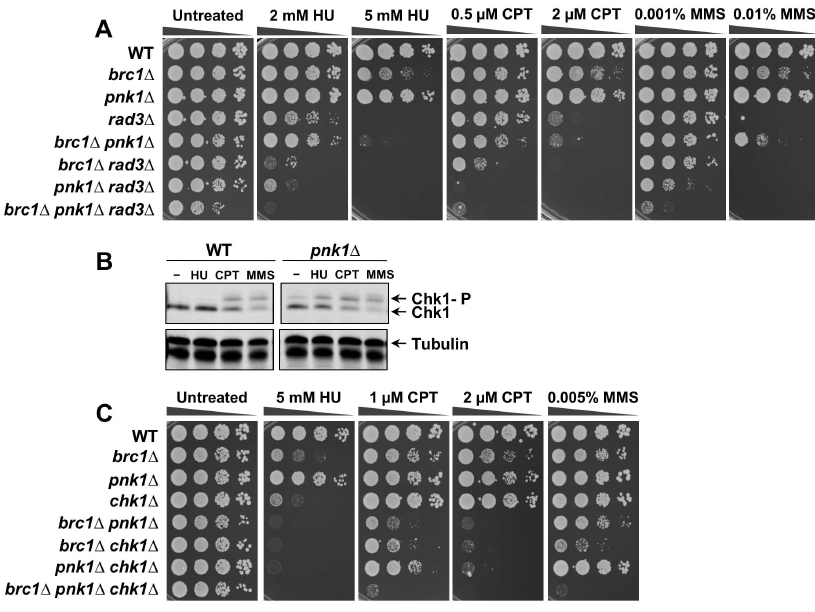
DNA damage checkpoint activation and requirement for checkpoint proteins in *pnk1Δ* cells. **A)** Effects of combining *pnk1Δ* with *rad3Δ* or *brc1Δ* mutations. Tenfold serial dilutions of cells were exposed to the indicated DNA damaging agents. Plates were incubated at 30°C for 3 to 4 days. **B)** After CPT and MMS treatment, Chk1 is phosphorylated in control cells, as indicated by the appearance of a slow-mobility species. Chk1 undergoes activating phosphorylation in untreated *pnk1Δ* cells. **C)** Effects of combining *pnk1Δ* with *chk1Δ* or *brc1Δ* mutations.

Rad3 phosphorylates the checkpoint kinase Chk1 in response to replication fork collapse (32-34). Immunoblot assays that detect phospho-Chk1 confirmed that Chk1 is activated even in the absence of genotoxin treatment in *pnk1Δ* cells (Figure 2B). No negative genetic interaction between *pnk1Δ* and *chk1Δ* was evident in the absence of genotoxins, indicating that the spontaneous DNA lesions causing Chk1 activation in *pnk1Δ* cells are efficiently repaired in the time frame of a normal G2 phase (Figure 2C, untreated panel). However, genotoxin treatment revealed a synergistic negative genetic interaction between *pnk1Δ* and *chk1Δ* that was most evident in cells treated with CPT. Chk1 was also critical in *pnk1Δ* cells treated with HU, which was consistent with the enhanced Chk1 phosphorylation in HU-treated *pnk1Δ* cells (Figure 2B). Elimination of Chk1 also enhanced the CPT and MMS sensitivity of *pnk1Δ brc1Δ* cells (Figure 2C).

From these results, we conclude that *pnk1Δ* cells accumulate DNA lesions that activate a Rad3-dependent checkpoint response leading to activation of Chk1. This response becomes especially critical when Brc1 is absent or when cells are treated with genotoxins that create SSBs.

### Increased frequency of Rad52 and RPA foci in *pnk1Δ* cells

These data indicated that *pnk1Δ* cells accumulate DNA lesions that activate DNA damage responses. To further test this proposition, we monitored foci formation of Rad52, which is normally essential for all forms of homology-directed repair in fission yeast. Mutants that suffer increased rates of replication fork collapse, or are unable to efficiently repair collapsed forks, typically display increased numbers of Rad52 nuclear foci (35-38). For these studies, we monitored Rad52 tagged with yellow fluorescent protein (Rad52-YFP) expressed from the endogenous locus. As observed previously (21), the frequency of cells with Rad52-YFP foci was significantly increased in *brc1Δ* cells (12.5%) compared to wild type (5.6%) (Figure 3A). The incidence of cells with Rad52-YFP foci was higher in the *pnk1Δ* strain (19%), and there was a further significant increase in the *brc1Δ pnk1Δ* strain (35.1%) (Figure 3A). Cell cycle phase analysis indicated that in all strains most of the cells with Rad52-YFP foci were in S-phase or early G2 phase, which suggests fork collapse as a primary source of these lesions. It was noteworthy that there was a large increase in mid- to late-G2 phase cells with Rad52 foci in the *brc1Δ pnk1Δ* strain (8.1%) compared to either single mutant (3.2% or 2.5%), respectively (Figure 3A). These data suggest Brc1 is required to efficiently repair lesions that accumulate in *pnk1Δ* cells, which could explain why Rad3 and Chk1 are crucial in *pnk1Δ brc1Δ* cells (Figure 2).

**Figure 3.**
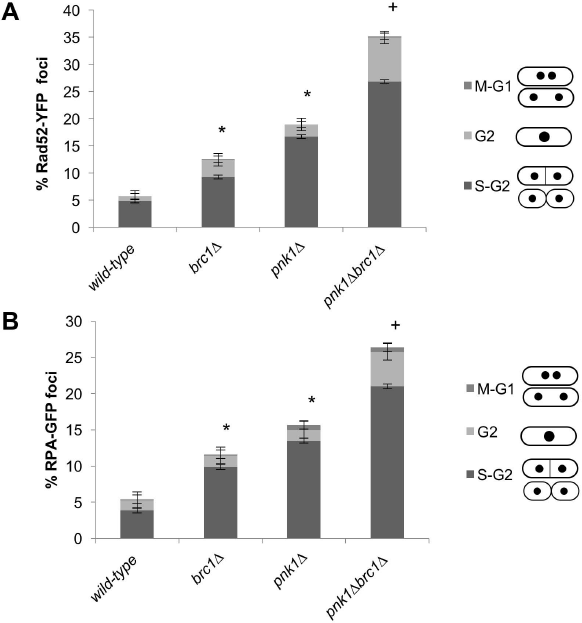
Rad52 and RPA foci increase in *pnk1Δ* cells and further increase in *brc1Δ pnk1Δ* cells. Cells expressing Rad52-YFP (A) or RPA-GFP (B) were cultured in minimal medium at 25°C until mid-log phase. Error bars correspond to standard deviation of the mean. Asterisk (*) and plus (+) symbols indicate statistically significant differences with wild type or *pnk1Δ* strains, respectively, as determined by two-tailed Student T-test, p-value ≤ 0.05.

In a separate experiment, we assessed foci formation by RPA, which is the major single-stranded DNA binding activity in eukaryotes. For these studies, we used strains that expressed the largest subunit of RPA, known as Ssb1 or Rad11/Rpa1, with a green fluorescent protein (GFP) tag (Figure 3B). The frequency of RPA-GFP foci was moderately increased in *brc1Δ* (11.6% versus 5.5% in wild type), further increased in *pnk1Δ* cells (15.7%), and even further increased in *brc1Δ pnk1Δ* (26.4%). As seen for Rad52, in all strains the RPA foci were predominantly observed in cells that were in S or early G2 phase, although combining the *brc1Δ* and *pnk1Δ* mutations did result in a substantial increase in mid- or late-G2 phase cells with RPA foci (4.8%), versus 1.6% in *brc1Δ* cells or 1.5% in *pnk1Δ* cells (Figure 3B).

These results are consistent with the synergistic growth defect and genotoxin sensitivity observed in *brc1Δ pnk1Δ* cells, and suggest that efficient repair of unligatable SSBs that accumulate in the absence of Pnk1 depends on Brc1.

### Mre11 and Ctp1 are crucial in the absence of Pnk1

Our studies suggested that lingering SSBs in *pnk1Δ* cells are converted to DSBs that require Brc1 for efficient repair. To further investigate this possibility, we assessed the requirements for the two major DNA end-binding protein complexes in fission yeast.

The Ku70/Ku80 heterodimer has a high affinity for DSBs. It promotes nonhomologous end-joining (NHEJ), which is critical for DSB repair in G1 phase when cells lack sister chromatids required for HDR (39). The *pku80Δ* mutation did not impair the growth of *pnk1Δ* cells or increase their sensitivity to UV, HU, CPT or MMS (Figure 4). These findings show that NHEJ does not play a significant role in an alternative pathway for repairing SSBs in the absence of PNKP.

**Figure 4.**
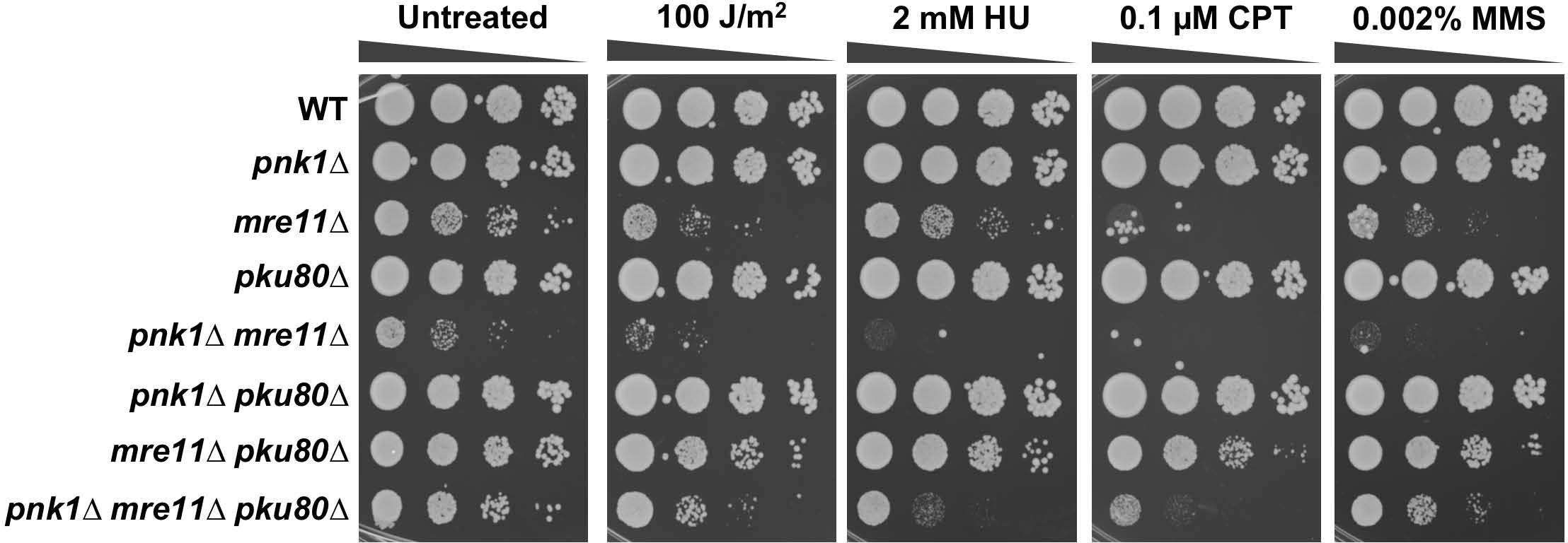
Mre11 is crucial in *pnk1Δ* cells. Effects of eliminating Mre11 or Pku80 in *pnk1Δ* background. Note that eliminating Ku partially suppresses the poor growth of *pnk1Δ mre11Δ* cells, but these cells remain acutely sensitive to the genotoxins. Tenfold serial dilutions of cells were exposed to the indicated DNA damaging agents. Plates were incubated at 30°C for 3 to 4 days.

The Mre11-Rad50-Nbs1 (MRN) endonuclease complex also binds DSBs, whereupon it endonucleolytically liberates Ku and initiates 5’-3’ resection to generate ssDNA tails required for HDR (40). These activities depend on Ctp1 (CtIP/Sae2), which is only expressed in S and G2 phases in *S. pombe* (41, 42). E-MAP studies indicated that both Mre11 and Ctp1 are likely to be important in the absence of Pnk1 (30, 43). Indeed, we found that *pnk1Δ mre11Δ* and *pnk1Δ ctp1Δ* double mutants grew very poorly compared to the respective single mutants (Figures 4 and 5). These negative genetic interactions were accentuated by exposure to DNA damaging agents (Figures 4 and 5).

**Figure 5.**
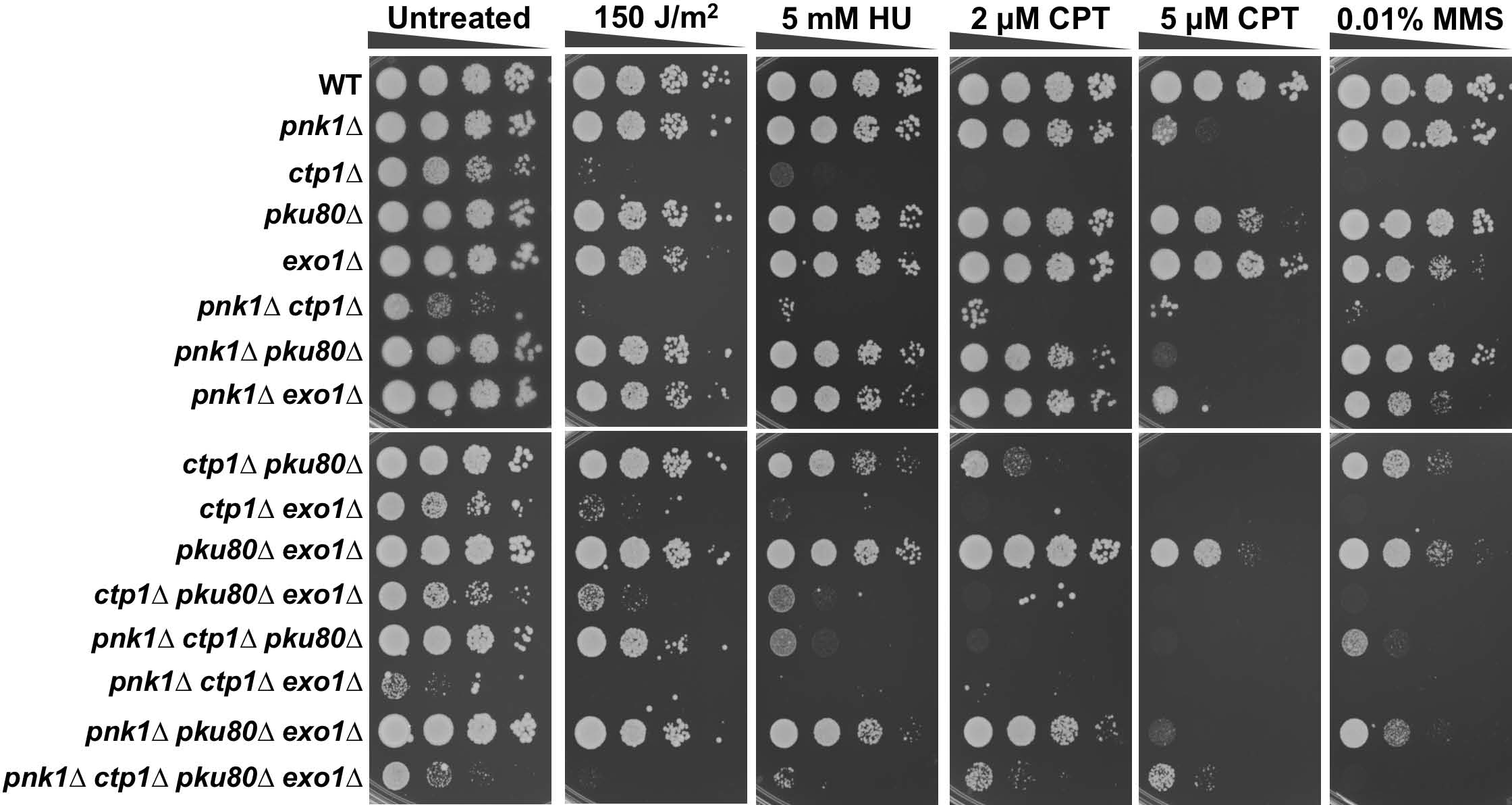
Ctp1 is critical in *pnk1Δ* cells. Note that eliminating Ku partially suppresses the poor growth of *pnk1Δ ctp1Δ* cells and this suppression requires Exo1. However, *pnk1Δ ctp1Δ pku80Δ* cells remain acutely sensitive to the genotoxins. Tenfold serial dilutions of cells were exposed to the indicated DNA damaging agents. Plates were incubated at 30°C for 3 to 4 days.

### Acute DNA damage sensitivity of *pnk1Δ mre11Δ pku80Δ* cells

The requirement for MRN and Ctp1 to initiate resection of DSBs can be substantially alleviated by genetically eliminating Ku, which allows Exo1 exonuclease to access DSBs and initiate resection (41, 42, 44). To investigate whether Exo1 effectivity substitutes for MRN-Ctp1 in the absence of Ku, we introduced the *pku80Δ* mutation into *pnk1Δ mre11Δ* and *pnk1Δ ctp1Δ* backgrounds. This analysis revealed that eliminating Ku partially restored growth and genotoxin resistance in these genetic backgrounds (Figures 4 and 5). In the case of *pnk1Δ ctp1Δ* cells, we confirmed that this suppression by *pku80Δ* depended on the presence of Exo1 (Figure 5). However, *pnk1Δ mre11Δ pku80Δ* cells grew poorly compared to *mre11Δ pku80Δ* cells and they were much more sensitive to the genotoxins (Figure 4). A similar effect was observed in the genetic studies involving *ctp1Δ* (Figure 5). These findings indicate that Exo1 has only a limited ability to substitute for MRN and Ctp1 in *pnk1Δ* cells.

### Spontaneous SSBs in *pnk1Δ* cells are repaired by Rad52-dependent HDR that does not require Rad51

The genetic requirements for Mre11 and Ctp1 strongly suggested that HDR resets replication forks that collapse at lingering SSBs in *pnk1Δ* cells. However, the critical HDR recombinase Rad51 was reported to be nonessential in these cells (14). We investigated these seemingly contradictory findings and confirmed that *pnk1Δ rad51Δ* cells grew nearly as well as *rad51Δ* cells (Figure 6A). The *pnk1Δ rad51Δ* cells were, however, more sensitive to several genotoxins, notably HU and MMS. These findings suggest that most of the spontaneous SSBs with 3’ phosphate that accumulate in *pnk1Δ* cells are repaired by an MRN-Ctp1-dependent mechanism that does not require Rad51.

**Figure 6.**
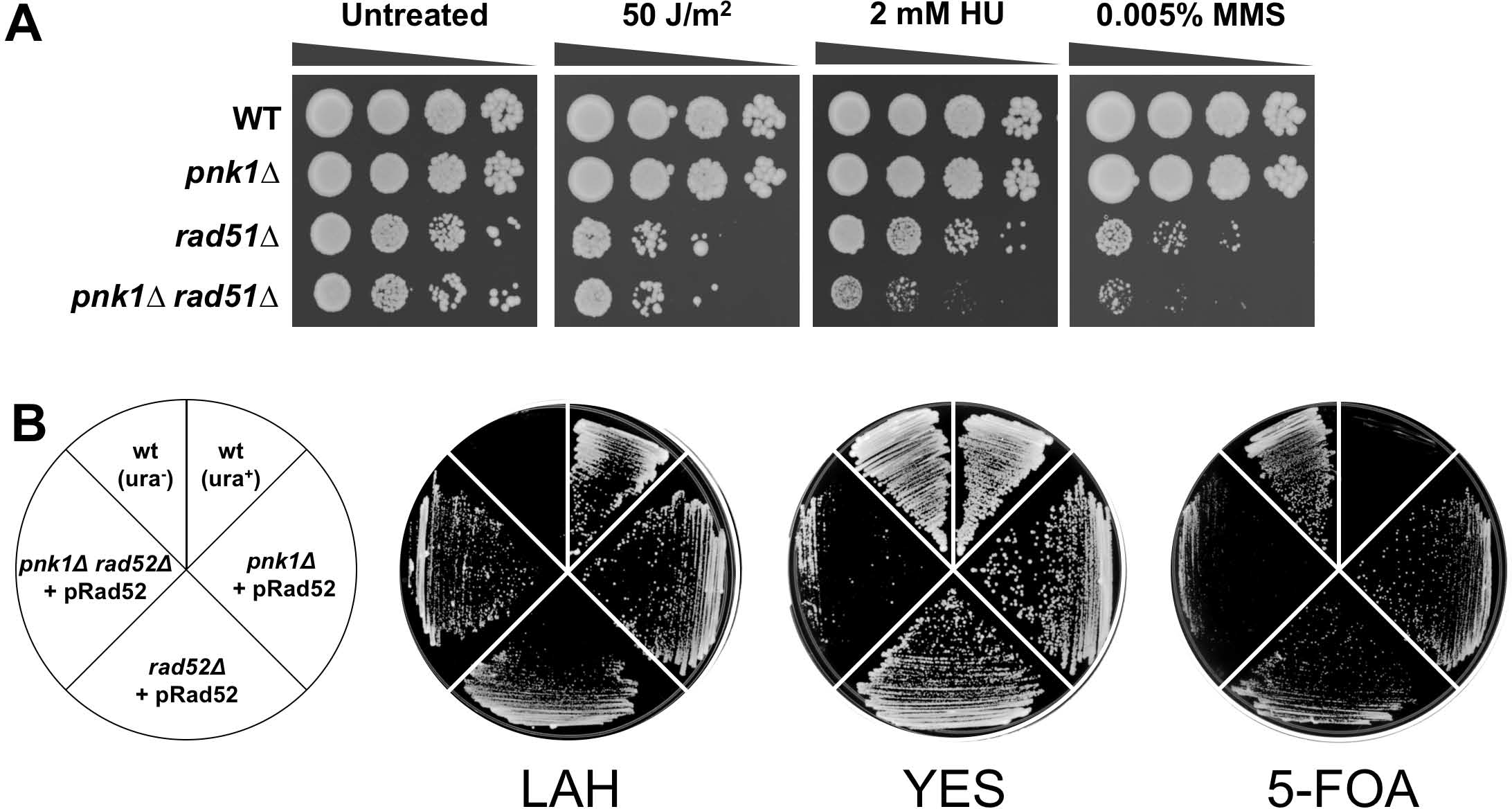
Requirement for Rad52 in *pnk1Δ* cells. **A)** A *pnk1Δ rad51Δ* is viable but it displays increased HU and MMS sensitivity relative to *rad51Δ.* Tenfold serial dilutions of cells were exposed to the indicated DNA damaging agents. Plates were incubated at 30°C for 3 to 4 days. **B)** Rad52 is crucial for viability in *pnk1Δ* cells. Strains with *pnk1Δ* or *rad52Δ* mutations, or the double mutant, in a *ura4-D18* background, were transformed with the pRad52 plasmid containing the *rad52^+^* gene and *ura4^+^* selectable marker. These strains and controls (wild type with *ura4^+^* or *ura4-D18*) were incubated on rich YES plates (no selection for *ura4^+^*), LAH media (selection for *ura4^+^*), or 5-FOA plates (counter selection for *ura4^+^*). Relative to *pnk1Δ* or *rad52Δ* single mutants carrying pRad52, the *pnk1Δ rad52Δ,* double mutant grew very poorly on 5-FOA plates, showing the Rad52 activity was crucial in the absence of Pnk1.

Previously, Whitby and co-workers reported that ~50% of CPT-induced collapsed replication forks are repaired by a Rad51-independent mechanism of HDR that requires Rad52 (45). Similarly, we found that elimination of the Swi1-Swi3 replication fork protection complex leads to collapse of replication forks that are repaired by a mechanism requiring Rad52 but not Rad51 (46). We set out to test whether Rad52 is critical in *pnk1Δ* cells. Genetic crosses involving *rad52Δ* are complicated by the frequent appearance of suppressors caused by loss of the F-box helicase Fbh1 (47). Therefore, we generated *pnk1Δ rad52Δ* or *rad52Δ* cells that were complemented by a pRad52 plasmid containing *rad52^+^* and the *ura4^+^* selectable marker (Figure 6B). Both strains grew relatively well in LAH medium that selects for the *ura4^+^* marker, but the *pnk1Δ rad52Δ* cells grew much more poorly in 5-FOA media that counter-selects against the *ura4^+^* marker. These data show that Rad52 is critical for cell viability in the *pnk1Δ* background. These results establish that many of accumulated spontaneous SSBs in *pnk1Δ* cells are repaired by a Rad52-dependent mechanism that does not require Rad51.

### Mus81-Eme1 Holliday junction resolvase and Rqh1 DNA helicase are essential in the absence of PNKP

In mitotic fission yeast, homology-directed repair of two-ended DSBs, for example as generated by ionizing radiation (IR), proceeds by synthesis-dependent strand annealing (SDSA). In SDSA, joint molecules do not mature into Holliday junctions, which explains why Mus81-Eme1 resolvase is not required for IR resistance in fission yeast (48-50). In contrast, HDR-mediated restoration of a broken replication fork produces a Holliday junction that must be resolved to allow chromosome segregation in mitosis, hence the acute requirement for Mus81 in conditions that increase replication fork collapse (45, 49, 51). We mated *pnk1Δ* and *mus81Δ* strains and found that the large majority of double mutant spores failed to yield viable colonies. The few viable double mutants were extremely sick (Figure 7A). We obtained the same results when we attempted to create *pnk1Δ eme1Δ* strains (Figure 7B).

**Figure 7.**
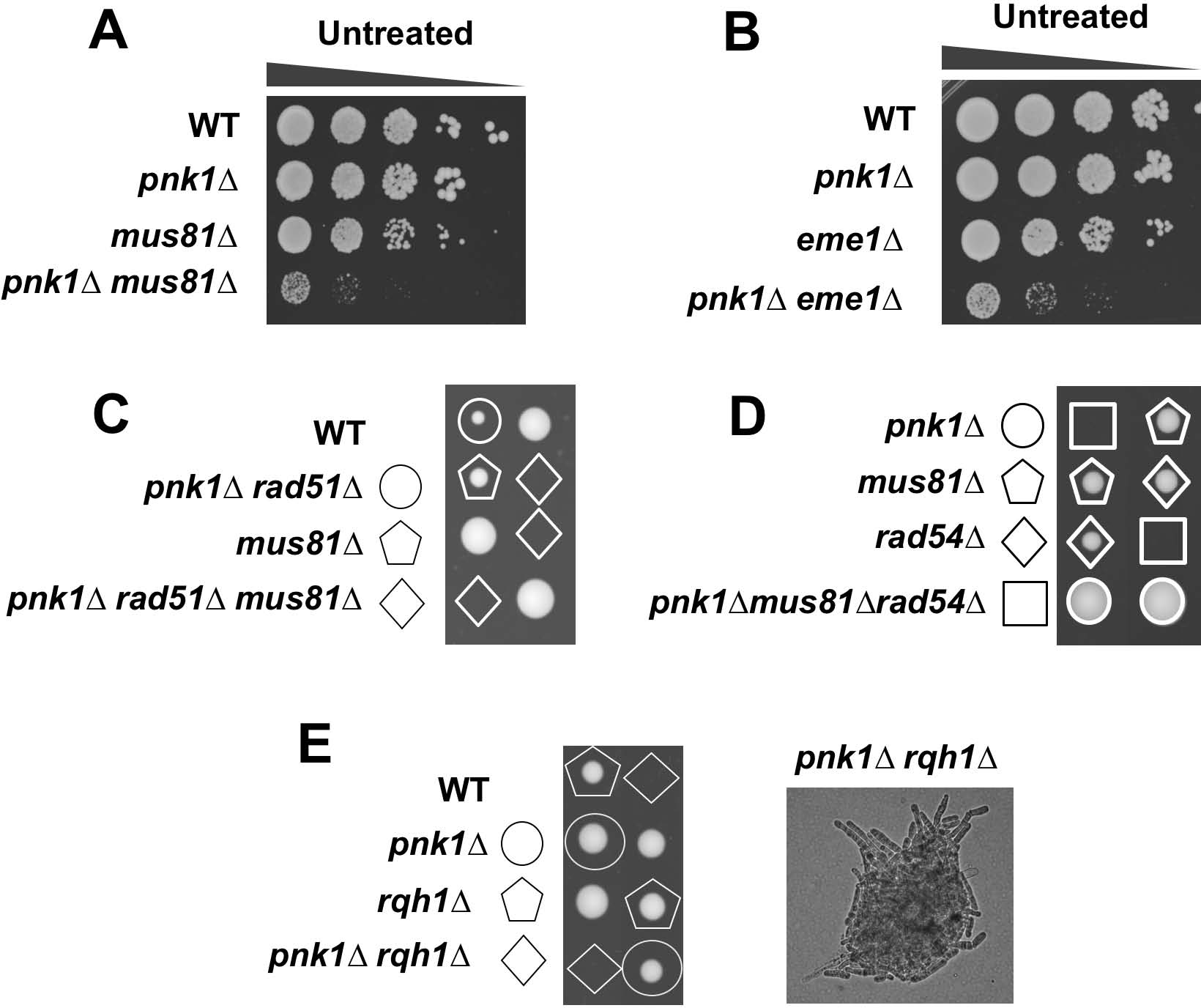
Mus81-Eme1 resolvase and Rqh1 DNA helicase are essential in *pnk1Δ* mutant cells. **A)** The few viable *pnk1Δ mus81Δ* viable cells recovered from genetic crosses are very sick compared to single mutants. **B)** The few viable *pnk1Δ eme1Δ* viable cells recovered from genetic crosses are very sick compared to single mutants. Tenfold serial dilutions of cells were plated and incubated at 30°C for 3 to 4 days. **C)** Elimination of Rad51 does not suppress *pnk1Δ mus81Δ* synthetic lethality. Tetrad analysis of *pnk1Δ rad51Δ* × *mus81Δ* cross. **D)** Elimination of Rad54 does not suppress *pnk1Δ mus81Δ* synthetic lethality. Tetrad analysis of *pnk1Δ rad54Δ* × *mus81Δ* cross. **E)** Tetrad analysis of mating between *pnk1Δ* and *rqh1Δ* strains. An example of germination products from a *pnk1Δ rqh1Δ* spore is presented in the right panel.

If HJs were formed without the participation of Rad51 in *pnk1Δ* cells, as suggested by our results, we would not expect loss of Rad51 to rescue the synthetic lethal interaction of *pnk1Δ* and *mus81Δ.* Indeed, genetic crosses showed that *rad51Δ* did not suppress *pnk1Δ mus81Δ* synthetic lethality (Figure 7C). We also investigated Rad54, which interacts with Rad51 and is required for Rad51-dependent HDR (52), but not the Rad52-dependent HDR of CPT-induced DNA damage that occurs independently of Rad51 (45). As predicted by our model, elimination of Rad54 failed to rescue the *pnk1Δ mus81Δ* synthetic lethality (Figure 7D).

Rqh1 is a RecQ family 3’-5’ DNA helicase that is orthologous to human WRN (Werner syndrome) and BLM (Bloom syndrome) DNA helicases, and *S. cerevisiae* Sgs1 DNA helicase (53, 54). Rqh1 is involved in multiple genome protection pathways and is particularly notable for its essential function in the absence of Mus81 (49). Strikingly, we found that Rqh1 is essential in the *pnk1Δ* background (Figure 7E).

### Swi10-Rad16 3’ flap endonuclease is not required in *pnk1Δ* cells

In *S. cerevisiae,* the viability of *tpp1Δ apn1Δ apn2Δ* cells depend on Rad10-Rad1 3’ flap endonuclease, which is orthologous to human ERCC1-XPF (11). These results suggest that Rad10-Rad1 provides an alternative mechanism for eliminating 3’-phosphates from DNA termini through endonucleolytic cleavage of 3’ DNA flaps. In fission yeast, *pnk1Δ apn2Δ* cells are inviable (14), but it remained possible that the 3’ flap endonuclease Swi10-Rad16 (55, 56), orthologous to budding yeast Rad10-Rad1, played an important role in repairing lingering SSBs in *pnk1Δ* cells. We found that *pnk1Δ swi10Δ* were viable and displayed no obvious growth defect relative to the respective single mutants (Figure 8). The *pnk1Δ swi10Δ* strain displayed slightly more sensitivity to HU and CPT but not MMS, but these genetic interactions did not appear to be synergistic. Thus, unlike key HDR proteins, Swi10-Rad16 3’ flap endonuclease is not part of a critical back-up mechanism for repairing SSBs with 3’ phosphate.

**Figure 8.**
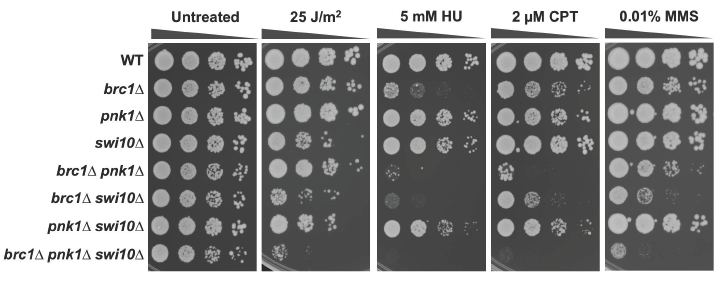
Swi10-Rad16 3’ flap endonuclease is not required in *pnk1Δ* cells. Tenfold serial dilutions of cells were exposed to the indicated DNA damaging agents. Plates were incubated at 30°C for 3 to 4 days.

### Increased spontaneous recombination in *pnk1Δ* cells

Finally, to explore whether Pnk1 deficiency creates perturbations to replication fork progression that increase recombination, we performed a mitotic intrachromosomal recombination assay. This assay determines the spontaneous frequency of Adenine positive (Ade^+^) colonies arising by recombination between two *ade6* heteroalleles flanking the *his3^+^* gene (57). Two classes of recombinants can be distinguished: deletion-types (Ade^+^ His^-^) and conversion-types (Ade^+^ His^+^) (Figure 9A). Total spontaneous recombination frequencies (deletion + conversion types, reported as events per 10^4^ cells) were increased ~3.4-fold in *pnk1Δ* cells (4.78 ± 1.16) compared with wild-type (1.41 ± 0.57). Although earlier studies indicated that spontaneous recombination frequencies in *brc1Δ* cells were strongly reduced (28), in our assays the spontaneous recombination frequencies of *brc1Δ* cells were not significantly different from wild type (Figure 9B). The spontaneous recombination frequency in *brc1Δ pnk1Δ* cells (3.91 ± 2.3) was moderately decreased compared to *pnk1Δ* cells (Figure 9B). Interestingly, conversion-type recombinants predominated in *pnk1Δ* cells. The statistically significant decrease of total recombinants in *brc1Δ pnk1Δ* cells compared to *pnk1Δ* was caused by a loss of conversion-type recombinants. Collectively, these data show indicate that a Pnk1 deficiency increases HDR-mediated genome instability.

**Figure 9.**
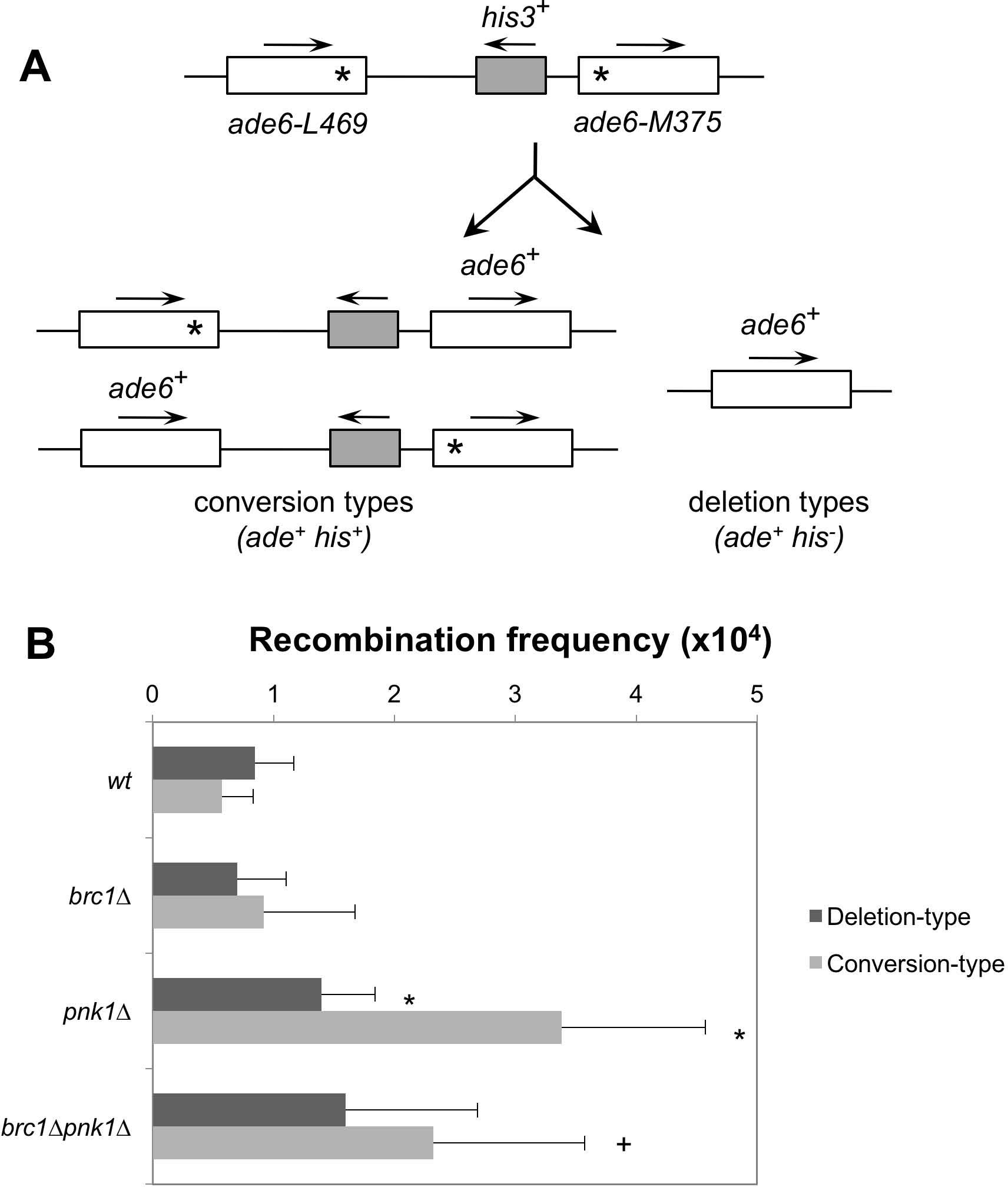
Increased spontaneous recombination in *pnk1Δ* cells. **A)** Schematic of the nontandem direct repeat of *ade6^-^* heteroalleles used for measuring spontaneous recombinant frequencies. Conversion types events result in Ade^+^ His^-^ colonies, whereas deletion types events result in Ade^+^ His^+^ colonies. **B)** Recombination frequencies (per 10^4^ viable cells ± SD) of the following strains: wild type (1.41 ± 0.57), *brc1Δ* (1.62 ± 1.16), *pnk1Δ* (4.78 ± 1.62), *brc1Δ pnk1Δ* (3.91 ± 2.3). Deletion types and conversion types were determined by replica-plating. Error bars correspond to standard deviations of the means. Asterisk depicts statistically significant differences with wild type and + symbol with *pnk1Δ,* as determined by two-tailed Student T-test, p-value ≤ 0.05.

## DISCUSSION

In this study, we have investigated how fission yeast cells tolerate the loss of polynucleotide kinase/phosphatase. In principle, a PNKP deficiency should result in lingering SSBs if there is no other efficient alternative mechanism for repairing SSBs with 3’ phosphate. Note that for this discussion we are presuming that genetic interactions involving *pnk1Δ* are caused by loss of 3’ phosphatase activity, as this defect is responsible for the DNA damage sensitivities of *pnk1Δ* cells, although it is formally possible that loss of 5’ kinase activity also contributes to these genetic interactions (14). SSBs can be converted into broken replication forks during S-phase. Broken forks are restored by homology-directed repair, thus key HDR proteins should be critical in absence of PNKP. Surprisingly, there is scant evidence on this point, and that which exists in fission yeast contradicts the model. Notably, the critical HDR protein Rad51 is not required in *pnk1Δ* mutants of fission yeast (14). We investigated this conundrum. Our experiments confirm that Rad51 is not essential in *pnk1Δ* cells; indeed, in the absence of exogenous genotoxins, *pnk1Δ rad51Δ* cells grow nearly as well *pnk1^+^ rad51Δ* cells. However, we have found that other HDR proteins become crucial for cell viability in the absence of Pnk1. Our studies established that Rad52 is essential in *pnk1Δ* cells. Similarly, *pnk1Δ mre11Δ* and *pnk1Δ ctp1Δ* strains are extremely sick. As the principal role of MRN complex and Ctp1 is to initiate resection of DSBs, these data strongly suggest that defective SSB repair in *pnk1Δ* cells is rescued by a mechanism that involves homology-directed repair of DSBs. Another key finding was the requirement for Mus81-Eme1 resolvase in *pnk1Δ* cells. As discussed above, Mus81-Eme1 is not required for survival of IR-induced DSBs, but it is crucial for recovery from replication fork breakage (49, 51, 58). Thus, our data strongly support the idea that lingering SSBs in *pnk1Δ* cells trigger replication fork collapse. This conclusion is further supported by the large increase in RPA and Rad52 foci in *pnk1Δ* cells, and cell cycle phase analysis indicating that most of the cells with these foci were in S-phase or early G2 phase.

The nature of the accumulating DNA lesions in *pnk1Δ* cells are also indicated by the negative genetic interaction with *brc1Δ*. As previously reported, *brc1Δ* cells are largely resistant to IR but quite sensitive to CPT, indicating that Brc1 functions in S-phase to assist the repair of collapsed replication forks (21). Thus, a defect in efficiently repairing collapsed replication forks most likely accounts for the synthetic sickness observed in *pnk1Δ brc1Δ* cells. Brc1 function partially depends on its ability to bind γH2A (21), hence it is noteworthy that mutations that specifically disrupt this binding show a synergistic negative genetic interaction with *pnk1Δ* when cells are exposed to CPT.

The absence of an obvious negative genetic interaction involving *pnk1Δ* and *chk1Δ* mutations in cells grown without genotoxins, despite the evident activation of Chk1, also provides clues about the DNA lesions that accumulate in the absence of PNKP. Chk1 delays the onset of mitosis by inhibiting Cdc25, which is the activator the cyclin-dependent kinase Cdc2 (59). Fission yeast has a naturally long G2 phase, thus activating a cell cycle checkpoint that delays mitosis should be less important if all DSBs are formed early in the cell cycle during S-phase. These facts explain why *chk1Δ* cells are relatively tolerant of moderate doses of genotoxins such as CPT, in which toxicity is mainly caused by breakage of replication forks, unless homology-directed repair is slowed by partial loss-of-function mutations in HDR proteins (60). These observations are consistent with a model in which replication forks break when they encounter lingering SSBs in *pnk1Δ* cells.

Neither *pnk1Δ* or *chk1Δ* mutants are strongly sensitive to 1 or 2 μM CPT, yet the *pnk1Δ chk1Δ* double mutant is acutely sensitive (Figure 2C). A similar genetic relationship is observed for *pnk1Δ* and *rad3Δ* (Figure 2A). These heightened requirements for checkpoint responses in *pnk1Δ* cells suggest that alternative repair pathways for repairing SSBs with 3’ phosphate are slow or inefficient. This interpretation is consistent with the increased level of Chk1 phosphorylation observed in the absence of genotoxin exposure in *pnk1Δ* cells (Figure 2B).

What is the explanation for the Rad51-independent repair of broken replication forks in *pnk1Δ* cells? As previously proposed, replication fork collapse caused by the replisome encountering a single-strand break or gap can generate a broken DNA end and a sister chromatid with a single-strand gap (45). This gap will tend to persist when it has a 3’ phosphate, as described below. The ssDNA gap may provide access to a DNA helicase that generates unwound donor duplex that participates in Rad52-mediated strand annealing (45). This process would not require Rad51. An alternative explanation concerns the location of lingering SSBs in *pnk1Δ* cells. In fission yeast, the ~150 copies of the ribosomal DNA locus are arranged in tandem repeats at each of chromosome III. We have previously reported that SIx1-SIx4 structure-specific endonuclease helps to maintain rDNA copy number by promoting HDR events during replication of the rDNA (61). Strikingly, these HDR events require Rad52 but not Rad51. Moreover, Mus81 and Rqh1 have crucial roles in maintaining rDNA in fission yeast (34, 62). If a large fraction of the lingering SSBs in *pnk1Δ* cells occur in the rDNA, this property could explain why Rad52, Mus81 and Rqh1 are required in *pnk1Δ* cells, but Rad51 is dispensable.

We note that whereas genetic elimination of Ku strongly suppresses the poor growth and genotoxin sensitivities of *mre11Δ* cells, which is explained by Exo1 resecting DSBs in the absence of Ku (40, 41), we observed that elimination of Pnk1 in *mre11Δ pku80Δ* cells impairs growth and greatly sensitizes cells to the DNA damaging agents. These data suggest that rescue of *mre11Δ* by Exo1 is inefficient in *pnk1Δ* cells. DSBs with 3’ phosphate might be poor substrates for Exo1, or they might attract other DNA end-binding proteins that block access to Exo1. MRN might resect 3’ phosphate ends at DSBs, thereby restoring 3’ hydroxyl (63). Alternatively, MRN might assist an alternative nuclease in repairing DSBs with 3’ phosphate. Human AP endonuclease efficiently removes 3’-PG at single-strand nicks but the corresponding activity at DSBs is very weak (64). Another possibility is that the acute genotoxin sensitivity of *mre11Δ pku80Δ pnk1Δ* cells simply reflects the fact that Pnk1 is required for a large fraction of the SSB repair in fission yeast, whether these SSBs arise from endogenous or exogenous DNA damaging agents. In this case, lack of Pnk1 will lead to a large increase of SSBs that are then converted to DSBs during replication, which might overwhelm the ability of Exo1 to substitute for MRN protein complex. Future experiments will be needed to test these models.

The 3’ phosphate responsible for the persistence of a SSB in *pnk1Δ* cells can itself be a barrier to the completion of homology-directed repair when the SSB is converted to a broken replication fork (65). Here, we consider models for tolerance of persistent SSBs with 3’ phosphate (Figure 10). When a replication fork collapses upon encountering a SSB with 3’ phosphate in the lagging strand template, the product is a one-ended DSB containing a 3’ phosphate (Figure 10, step 1a). Resection generates a single-strand overhang that invades the sister chromatid, but the 3’ phosphate blocks priming of DNA synthesis and restoration of an active replication fork (step 1b). This barrier to DNA synthesis might favor dissolution of the joint molecule, but resolution of the D-loop or Holliday junction by Mus81-Eme1 and ligase would stabilize the sister chromatid junction, allowing completion of replication by the converging fork (step 1c). The final product is a replicated chromosome containing a small ssDNA gap with the 3’phosphate (step 1d). When a replication fork collapses upon encountering a SSB with 3’phosphate in the leading strand template, the product is a one-ended DSB containing a 3’ hydroxyl opposite a sister chromatid with a ssDNA gap with 3’phosphate (Figure 10, step 2a). As previously noted (65), the SSB in the sister chromatid will block homology-directed repair, but replication by the converging fork will lead to replication fork collapse, leaving a DSB with a 3’phosphate (step 2b). At this point repair can proceed by SDSA (step 2c), eventually leading to one intact chromosome and the other containing a single-strand gap with a 3’phosphate (step 2d). Plans are underway to test these models.

**Figure 10.**
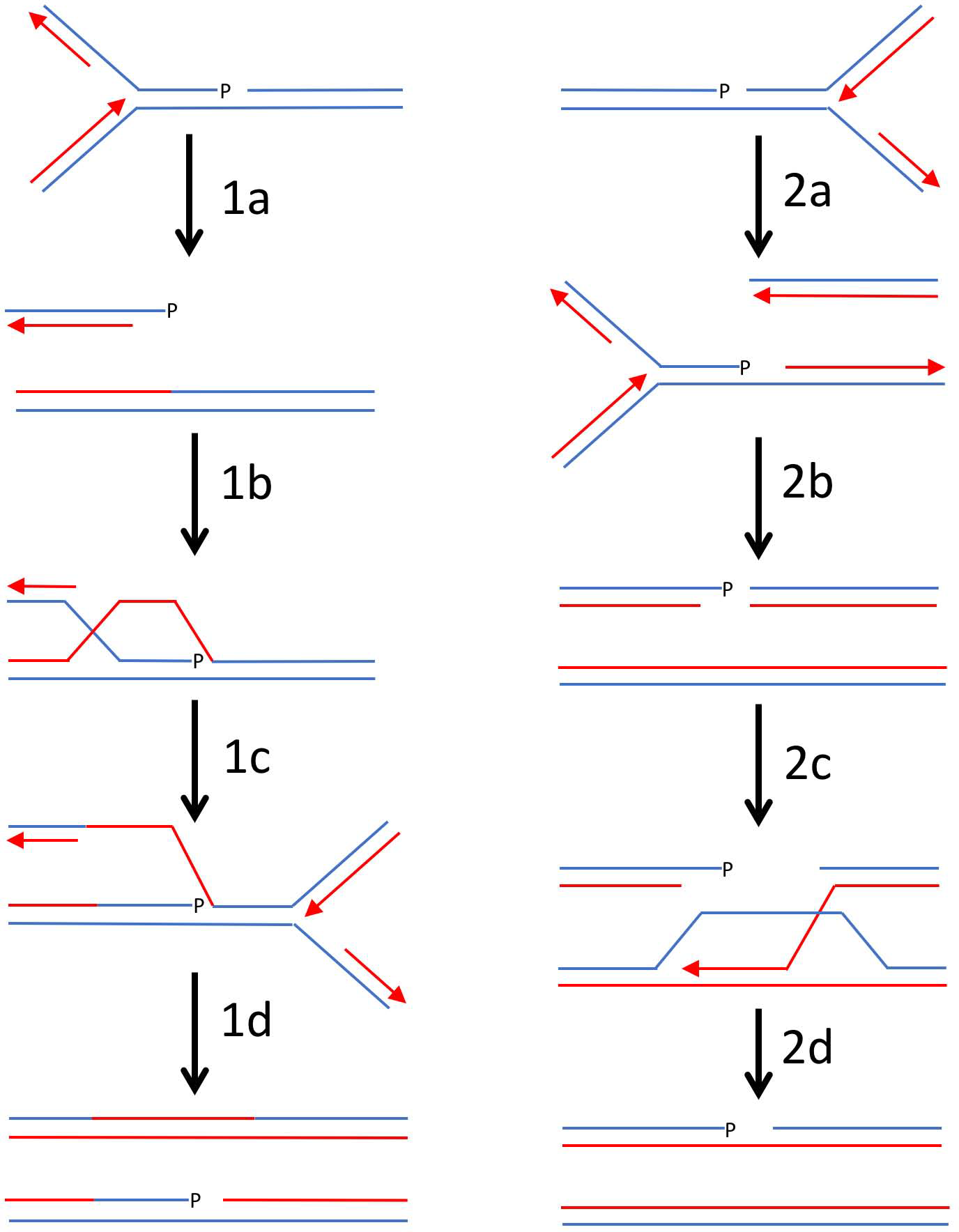
Models for replication-coupled repair of SSBs with 3’ phosphate terminus. 1a, replication fork collapses upon encountering a SSB with 3’ phosphate in the lagging strand template. 1b, resection of DSB followed by strand invasion of the sister chromatid using 3’ single-strand overhang containing 3’ phosphate. 1c, resolution of D-loop or Holliday junction by Mus81-Eme1. 1d, replication by converging fork, leaving a single-strand gap with 3’ phosphate. 2a, replication fork collapses upon encountering a SSB with 3’ phosphate in the leading strand template. 2b, converging fork collapses at SSB, leaving DSB with 3’ phosphate terminus. 2c, resection of DNA end with 3’ hydroxyl, followed by strand invasion of the sister chromatid. 2d, completion of repair by SDSA leaves a single-strand gap with 3’ phosphate.

In summary, these studies establish that polynucleotide/kinase phosphatase plays a crucial role in preventing the accumulation of SSBs that trigger replication fork collapse and genome instability in fission yeast, with the special property that many of these broken replication forks are repaired by an HDR mechanism that requires Mre11, Rad52 and Mus81, but not Rad51. With the recent evidence that Rad52 plays a crucial role in repair of broken replication forks in mammalian cells (66, 67), it will be of special interest to evaluate the importance of Rad52 in PNKP-deficient mammalian cells.

## MATERIALS AND METHODS

### Strains and genetic methods

The strains used in this study are listed in Table S1. Standard fission yeast methods were used (68). Deletion mutations strains were constructed as described (69). The *pnk1::KanMX6* strains were created from the wild-type strains using the PCR-based method and the primers, pnk1.G (5′-GTATGTTATTGAAACCACCCATTTTCATTGCTATGCAATTATAATATAGCTAACTCAATTACCAAGTCCCATTTAGTATT**CGGATCCCCGGGTTAATTAA**-3′) and pnk1.H (5′-ATAATTTTTATAAACGTTTGGTTTTAGTGGGATCAATAACTATATATTTTTGAAATTAATGCAATTTAATAATTTCTTAG **GAATTCGAGCTCGTTTAAAC**-3′). The nucleotide sequences in boldface overlap to the KanMX cassette of plasmid pFA6a-kanMX4. Successful deletion of these genes was verified by PCR. Tetrad analysis was performed to construct double mutants and verified by PCR.

### Survival assays

DNA damage sensitivity assays were performed by spotting 10-fold serial dilutions of exponentially growing cells onto yeast extract with glucose and supplements (YES) plates, and treated with indicated amounts of hydroxyurea (HU), camptothecin (CPT), and methyl methanesulfonate (MMS). For UV treatment, cells were serially diluted onto YES plates and irradiated using a Stratagene Stratalinker UV source. Cell survival was determined after 3-4 days at 30°C.

### Immunoblots

For Chk1 shift, whole cells extracts were prepared from exponentially growing cells in standard NP-40 lysis buffer. Protein amounting to ~100 mg was resolved by SDS-PAGE using 10% gels with acrylamide:bis-acrylamide ratio of 99:1. Proteins were transferred to nitrocellulose membranes, blocked with 5% milk in TBST (137 mM Sodium Chloride, 20 mM Tris, pH 7.6, 0.05% Tween-20) and probed with anti-HA (12C5) antibody (Roche).

### Microscopy

Cells were photographed using a Nikon Eclipse E800 microscope equipped with a Photometrics Quantix charge-coupled device (CCD) camera and IPlab Spectrum software. All fusion proteins were expressed at their own genomic locus. Rad52-yellow fluorescence protein (YFP) expressing strains were grown in EMM until mid-log phase for focus quantification assays. Quantification was performed by scoring 500 or more nuclei from three independent experiments.

### Recombination assay

Mitotic recombination was assayed by the recovery of Ade+ recombinants from the strains containing the intrachromosomal recombination substrate. Spontaneous recombinant frequencies were measured as described (57). Frequencies of fifteen colonies were averaged to determine the mean recombination frequency. Error bars indicate standard deviation from the mean. Two sample t-test were used to determine the statistical significance of differences in recombination frequencies.

## ACKNOWLEDGEMNTS

We thank Nick Boddy and Mariana Gadaleta for helpful discussions.

## SUPPORTING INFORMATION LEGENDS

**Table S1**. *S. pombe* strains used in this study.

